# Expanding the scope of redox-balance growth coupling techniques with a carbon cofeeding strategy

**DOI:** 10.64898/2026.04.01.713023

**Authors:** Aidan E. Cowan, Bridgie Cawthon, Mason Hillers, Sean Perea, Maxwell Grabovac, Anna Stanton, Samer Saleh, Jennifer Gin, Yan Chen, Christopher J. Petzold, Jay D. Keasling

## Abstract

Metabolic engineering to produce molecules not naturally synthesized by the host often requires directed evolution to improve pathway enzyme performance. Growth-coupled selection can dramatically increase directed-evolution throughput, and manipulation of redox balance has proven effective for tying reductase fitness to microbial growth. However, most redox-balance selections require feeding the reductase substrate because of stoichiometric constraints. This is impractical for many biosynthetic pathways either due to practical limitations on cost or complexity of bulk substrate synthesis, or the lack of an ability to transport substrate into cells, for example intracellular acyl-CoA/ACP intermediates. Here we define stoichiometric constraints that make substrate feeding necessary for many acetyl-CoA–derived reduction pathways in NADPH-imbalanced hosts. We overcome these constraints with a dual-feedstock strategy in which glucose provides reducing power while acetate supplies additional acetyl-CoA without directly perturbing redox balance. In an engineered *E. coli* selection strain, acetate co-feeding enabled growth coupling of acetaldehyde, 3-hydroxybutyrate, and mevalonate production and produced a linear correlation between product formation and growth. We then used this selection to evolve a class II HMG-CoA reductase (HMGR) from *Delftia acidovorans* toward NADPH utilization, enriching variants with improved NADPH-dependent activity. Finally, propionate co-feeding enabled growth coupling of propionyl-CoA reduction, supporting the generality of carbon co-feeding for selecting enzymes in pathways involving acyl-chain elongation and reduction.

**Highlights:**

- Stoichiometric limits of redox-balance growth coupling are defined
- Acetate co-feeding supplies acetyl-CoA without perturbing redox balance
- Co-feeding enables growth coupling of acetaldehyde, 3-HB, and mevalonate
- Growth coupling enables evolution of HMGR toward NADPH specificity
- Propionate co-feeding extends growth coupling to additional acyl-CoA substrates

## 1. Introduction

Microbial bioproduction offers tremendous potential to displace petroleum-based chemical engineering methods in the synthesis of chemicals, fuels, and materials.^1–3^ As a more sustainable alternative to petrochemical production, microbial bioprocesses convert renewable biological feedstocks into value-added products with a lower carbon footprint.^4,5^ Several bioproducts have achieved large-scale commercial production, including the natural fermentation products ethanol^6^ and lactic acid^7^.

Metabolic engineering has enabled the production of myriad other chemicals of industrial relevance. However, bioproducts produced through synthetic or heterologous pathways have garnered only modest market penetration in comparison.^2,8^ A major barrier to achieving a wider penetration of biomolecules into chemical markets is the production metrics—titers, rates and yields (TRY)—of many bioproduction pathways.^9,10^ A key tool for improving pathway flux is the directed evolution of bioproduction pathway enzymes. Directed evolution can be a powerful tool to adapt enzymes to the use of non-native cofactors or substrates to increase TRY of synthetic or unnatural pathways.

Directed evolution is an iterative process of mutagenesis and screening to enhance protein function, and has proven to be a powerful strategy for engineering biological catalysts with novel or enhanced activities.^11^ This approach has been instrumental in expanding the functional repertoire of enzymes beyond their natural capabilities and holds immense promise for enabling high-TRY bioproduction of a greater diversity of synthetic pathways.^12^ However, classical directed evolution approaches for enzymes are limited by their screening capabilities, often requiring an individual assessment of each mutant of interest. Techniques have emerged to greatly increase the throughput of evaluating individual enzyme sequences, however these techniques are not universally applicable to metabolic enzymes, oftentimes requiring a colorimetric or fluorometric output.^13–15^ Regardless, despite major advances in high-throughput enzyme screening, experimental evaluation remains limited relative to the vastness of sequence space (∼20^100^ variants for a 100–amino-acid protein) and is still labor- and cost-intensive.^16,17^

To address this constraint, growth-coupled selection mimics natural selection by tying protein fitness to host survival, greatly increasing the throughput of DNA library evaluation.^18–22^ The only limitation to the number of variants which can theoretically be assayed using growth-coupled selection is the transformation efficiency of a given DNA library or the culture volume used for selection.^21,23^ By coupling enzyme function to cellular fitness, growth-coupled selection approximates natural selection in the laboratory and enables DNA library evaluation at scales that approach the practical limits of sequence space exploration. However, there is no universal selection system for evolving protein function and developing a robust selection strategy for a given enzyme can prove to be more laborious and challenging than simply relying upon established screening methods. Developing a more generalizable approach to growth coupling of metabolic enzymes is a core goal of this work.

The development of a generalizable selection system for enzyme or pathway activity, one that leverages a shared feature among diverse bioproduction pathways, could significantly enhance our capacity to engineer high-TRY bioproduction to replace petrochemicals. Many petroleum-derived chemicals, including fuels and plastics, are significantly more reduced than central metabolic intermediates or biological feedstocks like glucose, a characteristic that reflects their origin from the highly reduced hydrocarbons found in crude oil.^24^ To synthesize highly reduced products from metabolic intermediates and sugar-based feedstocks, cells must utilize pathways involving oxidoreductases and their associated redox cofactors to facilitate the necessary electron transfer reactions.^24,25^ Selection schemes exploiting the activity of these enzymes would therefore be broadly applicable to a variety of biochemical pathways capable of producing petrochemical substitutes such as the polyketide^26–29^, isoprenoid^30^ and fatty-acyl^31^ production pathways. Growth-coupled oxidoreductase activity, where the rate of cell division depends entirely upon the activity of a specific set of oxidoreductases, is a natural characteristic of anaerobic fermentation. Anaerobically, when grown on substrates such as glucose, microorganisms generate fermentation products such as ethanol, lactic acid, and succinate in order to regenerate NAD^+^ as an electron acceptor for glycolysis. This selective pressure has, over evolutionary time, evolved these pathways to become incredibly efficient, a characteristic which industrial biotechnology has exploited to achieve unrivaled penetration of these products into chemical markets.

Engineering selection systems which rely upon redox balance to select for the activity of oxidoreductases within growing populations of cells have become an attractive strategy for high-throughput directed and adaptive evolution.^21^ Anaerobically grown *Escherichia coli* deficient in fermentation pathways, and therefore unable to regenerate NAD^+^, can be rescued by the activity of reductase enzymes.^32^ Partial fermentation deficits have been used to drive the redirection of metabolic flux towards specific native fermentation products to enhance titer and yield.^33,34^ Heterologously expressed fermentation pathways, which are sufficiently reducing to balance redox demands, have been used to rescue the growth of fermentation-deficient *E. coli*, generating driving forces which have allowed for directed evolution and pathway expression tuning.^32,35–37^ In both aerobic and anaerobic contexts, engineered bacterial strains unable to regenerate oxidized electron acceptors in the form of NAD^+32,38,39^, NADP^+40–42^, or the orthogonal redox cofactor NMN^+34^ have been employed for the directed evolution of various oxidoreductase enzymes. However, most such efforts have been constrained to (i) highly reduced products that intrinsically satisfy redox stoichiometry (often native fermentation products), or (ii) systems in which the immediate substrate for the reduction step can be supplied exogenously and imported, thereby decoupling redox recycling from the pathway’s intrinsic stoichiometry.^39^ This limits the applicability of redox-balance-based growth-coupled selections, excluding enzymes whose substrates cannot be supplied in selection media because they are toxic at high concentrations, prohibitively expensive or difficult to synthesize, or not membrane-permeable (such as CoA- or ACP-bound intermediates).

In this work, we investigated the limitations of redox-balance-based selection systems for the directed evolution of reductases. We identified strict stoichiometric constraints that govern the bioenergetic feasibility of coupling enzyme activity to growth through NADP⁺ regeneration and thereby restrict the classes of pathways that can be growth-coupled through redox balance. To overcome these limitations, we implemented an aerobic dual-feedstock strategy using glucose as a primary carbon source and acetate as a redox-neutral substrate for a heterologous pathway incapable of supporting growth alone. This strategy resolves redox stoichiometric constraints and enables growth coupling for various pathways that partially reduce elongating acyl-CoA chains, a key reaction shared between many pathways of biotechnological interest. We demonstrate this selective pressure is sufficient for directed evolution by selecting for and identifying variants of a reductase enzyme with altered cofactor specificity.

## 2. Results

### 2.1. Growth coupling the partial reduction of various acetyl-CoA-derived acyl-CoAs

To growth couple partially reducing biosynthetic pathways, we chose to employ a strain deficient in NADP^+^ regeneration due to NADPH’s role as the key biosynthetic electron donor. We started with *E. coli* MX203, which contains deletions in genes *pgi* and *edd* that disrupt Entner-Doudoroff (ED) and Embden-Meyerhof-Parnas (EMP) glycolysis and thereby reroute glucose metabolism through the pentose phosphate pathway (PPP) (Fig. 1a).^40^ The PPP generates NADPH and additional deletions in *qor* and *sthA* inhibit electron flux from accumulated NADPH into the quinone and NADH pools, respectively (Fig. 1a).^40^ Taken together, these deletions generate a strain that, when grown on glucose, exhibits severe growth limitation due to an accumulation of NADPH with the inability to regenerate sufficient NADP^+^ to act as an electron acceptor for the PPP.^40^

**Fig. 1.**
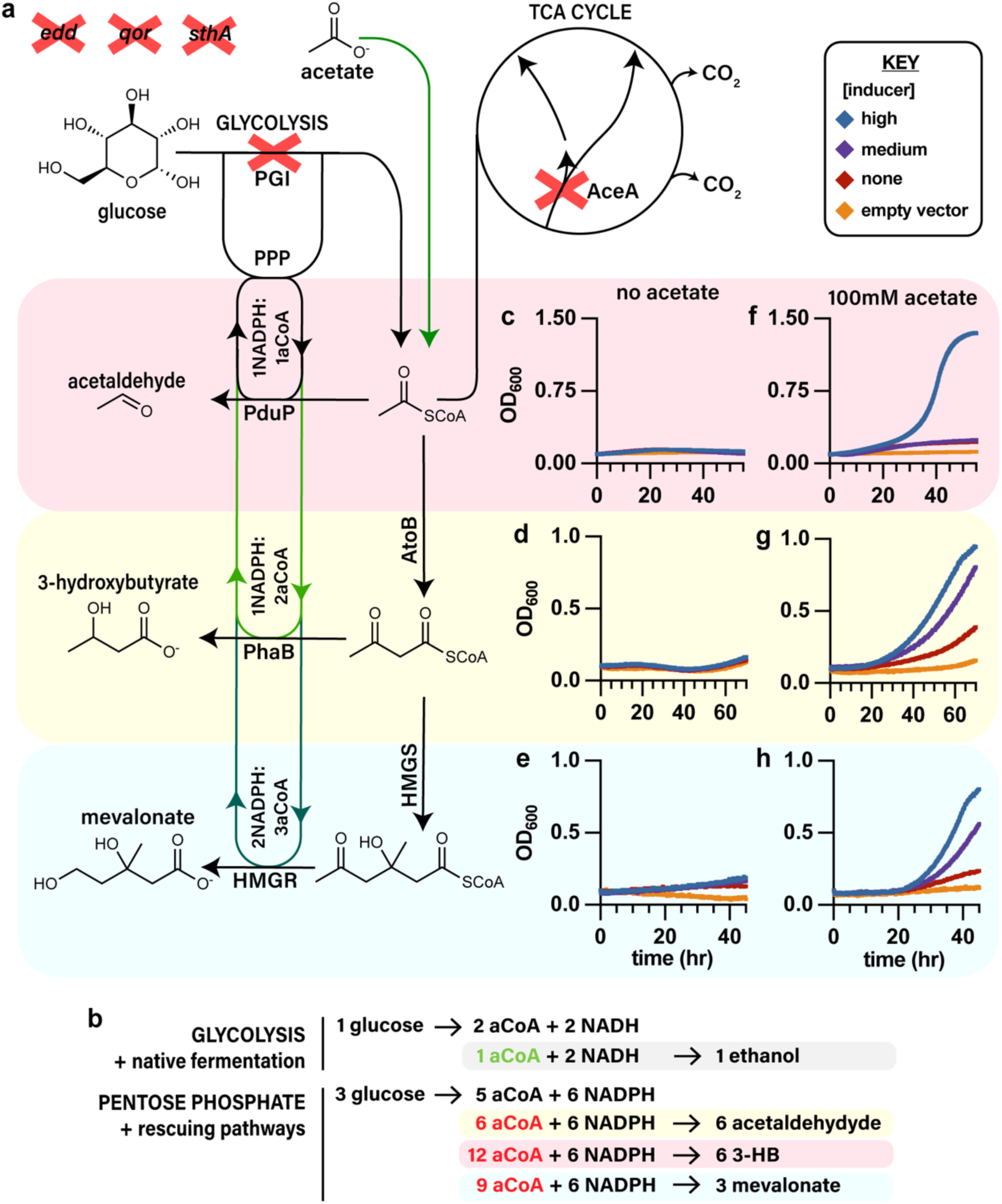
Growth coupling of partially reduced acyl-CoAs a, Schematic depicting the genetic modifications to enable the overaccumulation of NADPH (Δ*pgi*, Δ*edd*, Δ*qor*, Δ*sthA*) and the deletion of *aceA* to prohibit growth on acetate as a carbon source. Pathways enabling growth on a mixture of acetate and glucose are shown below and highlighted (red: acetaldehyde (strain APEQS_PduP), yellow: 3-HB (strain APEQS_3-HB), blue: mevalonate (strain APEQS_MEV_sa)). **b,** Simplified metabolic stoichiometries showing only bioavailable carbon and relevant reducing equivalents and assuming all carbon flows to acetyl-CoA (our pathways’ precursor molecule). Simplified stoichiometry of *E. coli* fermentative metabolism and APEQS metabolism when grown on glucose, showing redox-balanced fermentative or rescue pathways below. Ethanol fermentation requires less acetyl-CoA (green) than is produced from one glucose when redox balanced, reflecting its suitability as a fermentation pathway. Partially reducing pathways consume more acetyl-CoA (red) than is made available per unit glucose when redox balanced, indicating their inability to resolve redox balance in APEQS without acetate co-feeding. **c,** Unsuccessful growth coupling of strain APEQS_PduP in the absence of acetate (n=3). **d,** Unsuccessful growth coupling of strain APEQS_3-HB in the absence of acetate (n=3). **e,** Unsuccessful growth coupling of strain APEQS_MEV_sa in the absence of acetate (n=3). **f,** Successful growth coupling of strain APEQS_PduP when the strain is grown with additional 100 mM sodium acetate to satisfy stoichiometric constraints (n=3). **g,** Successful growth coupling of strain APEQS_3-HB when the strain is grown with additional 100 mM sodium acetate to satisfy stoichiometric constraints (n=3). **h,** Successful growth coupling of strain APEQS_MEV_sa when the strain is grown with additional 100 mM sodium acetate to satisfy stoichiometric constraints (n=3). All experiments were conducted in a MOPS medium containing 2% glucose with or without 100 mM acetate. Various concentrations of IPTG were added to modulate the induction of the three partially reducing pathways (high [IPTG]: blue (0.5 mM for p15A-based A5c backbone; 0.05 mM for ColE1-based pQE backbone), medium [IPTG]: purple (0.05 mM for p15A-based A5c backbone; 0.005 mM for ColE1-based pQE backbone), no IPTG: red). An empty vector control was included to demonstrate growth without leaky expression (orange). All growth experiments were repeated a minimum of 3 times and showed identical results.

We attempted to rescue the growth of MX203 using a variety of pathways involving the partial reduction of an elongated acyl-CoA to generate acetaldehyde, 3-hydroxybutyrate (3-HB) and mevalonate in an NADPH-dependent manner (Fig. 1a). The partial reduction of acetyl-CoA to generate acetaldehyde was accomplished by expressing an NADPH-dependent variant^43^ of the propanediol utilization protein from *Rhodopseudomonas palustris* (*Rp*PduP-NP) which natively reduces propionyl-CoA to propionaldehyde and shows promiscuous activity with acetyl-CoA and other CoA-ester substrates.^44^ *Rp*PduP-NP was expressed by plasmid pQE_PduP. 3-HB was produced via a two-step pathway in which overexpressed *E. coli* thiolase AtoB condenses two acetyl-CoA molecules to form acetoacetyl-CoA, which is then reduced by *Cupriavidus necator* PhaB to 3-hydroxybutyryl-CoA. 3-hydroxybutyryl-CoA is cleaved by a native thioesterase in *E. coli* to generate 3-HB.^45^ *phaB* and *atoB* were expressed using plasmid pBbA5c_3-HB. The production of mevalonate was catalyzed by AtoB, as well as HMGS and HMGR from *Staphylococcus aureus* (HMGR_sa). These genes catalyze the generation of acetoacetyl-CoA, 3-hydroxy-3-methylglutaryl-CoA (HMG-CoA) and mevalonate, respectively and were expressed using plasmid pBbA5c_MEV_sa.

Despite consuming NADPH and regenerating the essential cofactor NADP⁺, none of these pathways restored growth of MX203, even though they are robustly expressed (Supplementary Fig. 1). This outcome reflects a fundamental stoichiometric constraint in growth-coupled selection strategies based on redox balance. During fermentation, a fraction of available carbon must be diverted to reduced end products that serve as electron sinks. For example, during ethanol fermentation in *E. coli*, one acetyl-CoA molecule can be reduced by two NADH molecules to produce ethanol (Fig. 1b). This fully oxidizes the two NADH generated by glycolysis and allows for rebalancing of the cellular redox state. Because only a portion of the bioavailable carbon is diverted to fermentation, the remaining carbon in the form of pyruvate or acetyl-CoA can be used for growth and division (Fig. 1b). However, because the pathways we are working with are less reducing per unit carbon than ethanol fermentation, they require the diversion of a larger amount of bioavailable carbon to fully regenerate the fixed amounts of NADPH being generated by the PPP. When one of these partially reducing pathways is employed to rebalance the redox state of MX203 we observed that the amount of acetyl-CoA generated by the PPP was insufficient to enable the total oxidation of NADPH generated through the PPP (Fig. 1b). In previous work, these stoichiometric constraints have been remedied by the feeding the direct substrate of the target enzyme.^32,39,46^ However, all of these partially reducing pathways act upon acyl-CoAs which cannot cross the cell membrane and are difficult and costly to synthesize.

To address these constraints and allow for growth coupling we devised a co-feeding strategy where glucose would remain the primary carbon source while acetate would be co-fed to generate acetyl-CoA in a redox-neutral manner. This would enable generation of the substrate for each of these pathways without inducing additional redox imbalance, thereby decoupling NADPH-producing central metabolism and NADPH-consuming biosynthetic pathways to satisfy stoichiometric constraints otherwise precluding successful growth coupling. However, we also needed to prevent the cell from adapting to instead grow on acetate, while maintaining its ability to activate acetate to acetyl-CoA. To accomplish this, we made single deletion in the gene *aceA*, isocitrate lyase, which catalyzes the first step of the glyoxylate shunt. Without the glyoxylate shunt, acetate metabolism must proceed through the oxidative TCA cycle, which contains two decarboxylative steps. This means that there is no net carbon assimilation as both carbons on acetate are lost as CO_2_.^47^ *aceA* was deleted from MX203 by allelic exchange and the resulting strain was renamed APEQS to reflect its five gene deletions (*aceA, pgi, edd, qor, sthA*) (Supplementary Table 1).

Plasmids pQE_PduP, pBbA5c_3-HB and pBbA5c_MEV_sa were transformed into APEQS, generating strains APEQS_PduP, APEQS_3-HB and APEQS_MEV_sa. These strains were then grown in a minimal medium containing 2% glucose with or without 100 mM acetate to evaluate the ability of the reducing pathways to rescue the growth of APEQS. As expected, in the absence of acetate the strains showed no indication of growth coupling (Fig. 1c-e). However, in the presence of 100 mM acetate, where stoichiometric constraints to growth coupling could be addressed by co-feeding, each of these pathways was able to rescue the growth of APEQS (Fig. 1f-h).

Furthermore, for each strain, growth exhibited a dose-dependent relationship with the degree of IPTG-mediated induction of the corresponding plasmid with the highest concentration of inducer exhibiting the fastest growth and with an empty vector control showing the slowest growth. These results indicate that product formation in each of these pathways was successfully linked to growth rate.

### 2.2. Defining the dynamic range of selection and addressing limitations to generalizing growth coupling

An often undefined or underdefined metric of growth-coupled selection platforms is the dynamic range of the growth-coupled phenotype. In this study, the dynamic range is defined as the range of titers that elicit a meaningful change in growth rate. Defining the linear range of selection is important to allow researchers and engineers to understand what pathways may be suitable for growth coupling through a given selection scheme. If titers are above the dynamic range, then additional gains of function may not elicit meaningful improvements to growth and therefore not be selected for. Similarly, if titers are below the dynamic range of selection, even large (e.g. tenfold) improvements in titer will only elicit minor, if any, change in growth rate, making selection difficult or impossible.

To define the linear range of selection, we quantified 3-HB and mevalonate titers produced by APEQS strains under growth-coupled conditions in selective medium. We assayed the titer produced once strains had grown to their terminal optical density (OD) under various induction strengths of their respective pathways. This allowed us to correlate titer to growth across a range of titers and growth rates. The final titer achieved at each concentration of IPTG was highly correlated with growth rate across both 3-HB and mevalonate data (Fig. 2a). To further improve the correlation and better integrate 3-HB and mevalonate data from pathways with different degrees of reduction per unit carbon, we calculated the moles of electron pairs transferred to HMG-CoA or acetoacetyl-CoA necessary to generate the observed titer of mevalonate (2 mol electron pairs/mol mevalonate) and 3-HB (1 mol electron pairs/mol 3-HB), respectively for each sample. This measure of extracellular electron pairs (EEP) in molar units was more highly correlated with growth rate due to the normalization of titer units for 3-HB and mevalonate for their respective oxidation states (Fig. 2b).

**Figure 2.**
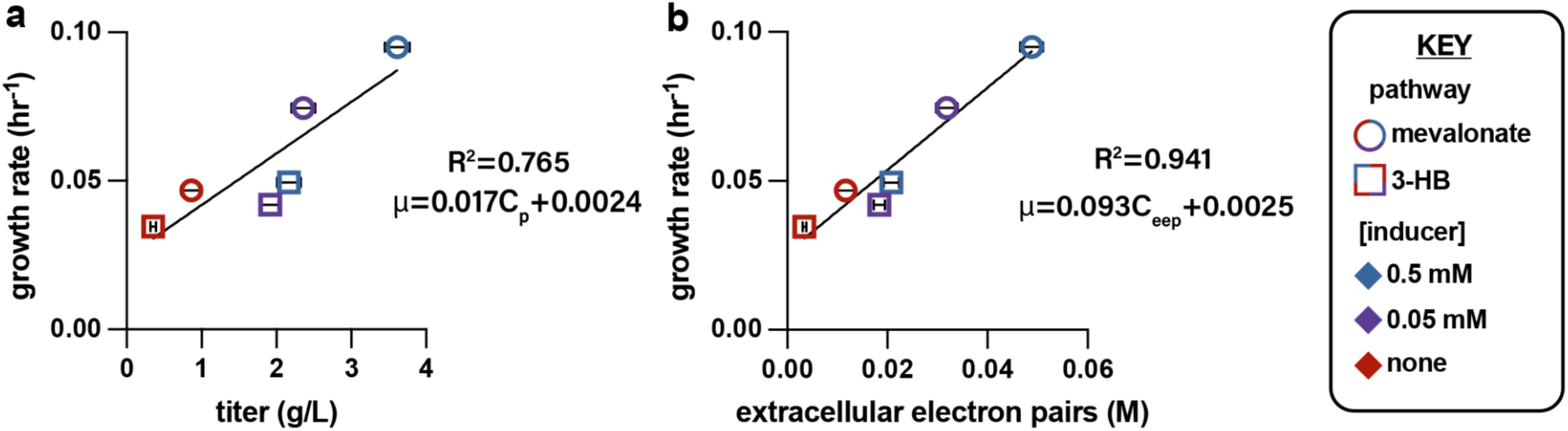
Growth rate and product formation correlation a, Final titer and growth rate were determined for 3 levels of induction of APEQS_MEV_sa and APEQS_3-HB and the correlation (R^2^ = 0.765) is shown along with a line of best fit (*C_p_*: concentration of product (titer g/L)) (n=3). The line of best fit was determined by the least-squares method. **b**, The extracellular electron pairs were calculated as the total number of electron pairs (arising from NADPH equivalents) required to produce a given titer from acetyl-CoA. This measure of extracellular electron pairs (EEP) in molar units (mol/L) allows for comparison of products with different oxidation states and improves the correlation between product formation and growth rate (R_2_ = 0.941) (n=3).

Across varying induction levels, product titer correlated linearly with growth rate for both the 3-HB and mevalonate rescue pathways (Fig. 2). Titers ranged from 0.35 ± 0.04 g/L 3-HB (0.0034 ± 0.0004 M EEP) to 3.6 ± 0.2 g/L mevalonate (0.049 ± 0.002 M EEP). Lines of best fit were determined using least-squares regression and are shown in Fig. 2. These lines of best fit can allow us to model how the selection might behave at low titers, and could help estimate if growth coupling is feasible for a given acyl-CoA reducing pathway using APEQS. For example, a 10-fold improvement in titer from 5 mg/L to 50 mg/L of mevalonate would only elicit an approximately 1% difference in growth rate, which is likely not sufficient to be selected for in a population of APEQS cells given the inherent variability between measurements. This semi-quantitative approach can give protein engineers a framework to understand the suitability of APEQS or related strains for their given engineering goals and define a general range of titers necessary to establish growth coupling.

### 2.3. Growth-coupled evolution of NADH-dependent HMGR for NADPH specificity

As a demonstration of the utility of the described growth coupling strategy, we sought to perform directed evolution. For the purposes of a limited demonstration, we chose to evolve NADPH specificity starting with a NADH-specific variant of the HMGR enzyme responsible for catalyzing the four-electron reduction of HMG-CoA to mevalonate. We generated a pooled site-saturation library of the NADH-dependent HMGR protein from *Delftia acidovorans* (HMGR_da) ^48^ which contained randomized NNK codons at three positions (D146, V148, L152) predicted to determine cofactor specificity^49^ (Fig. 3a, Supplementary Fig. 2a). Plasmid pBbA5c_MEV_da, which also contained *E. coli atoB* and *Saccharomyces cerevisiae hmgs* in the same operon as HMGR_da, served as the template for the generation of this library. The DNA library, which contained a theoretical maximum of 8,000 mutants, was used to transform the APEQS strain in triplicate and over 80,000 transformants from each replicate were pooled to minimize loss of library diversity. As controls, the original template plasmid pBbA5c_MEV_da, pBbA5c_MEV_sa (expressing NADPH-dependent HMGR_sa), and an empty plasmid backbone (pBbA5c_empty) were also used to transform APEQS, generating over 80,000 transformants of strains APEQS_MEV_sa, APEQS_MEV_da and APEQS_MEV_empty, respectively.

**Figure 3.**
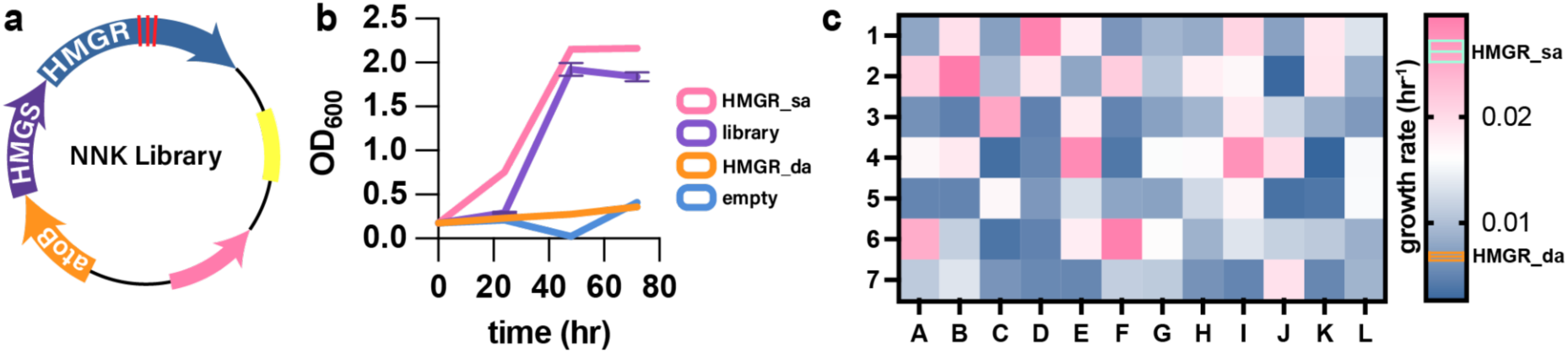
Growth-coupled evolution of NADH-dependent HMGR for NADPH specificity a, Schematic depicting HMGR_da library with three sites undergoing saturation mutagenesis highlighted by red lines. **b,** Growth of library and control cultures over the course of the HMGR_da evolution. APEQS was transformed with pBbA5c_MEV_sa (pink), library (purple), pBbA5c_MEV_da plasmid (orange) and pBbA5c_empty (blue) DNA. OD600 was measured periodically over the course of the 72-hour growth and evolution. **c,** Heat map showing the growth rate of 84 APEQS strains transformed with plasmids isolated after the evolution of HMGR_da. The growth rate of APEQS_MEV_sa and APEQS_MEV_da are indicated on the color key (right) by cyan and orange boxes respectively. The boxes represent one standard deviation from the mean value denoted as a horizontal line.

The library (in triplicate) and pooled control transformants were inoculated into selective medium containing glucose and acetate and cultured for 72 hours to enrich for HMGR_da variants with increased NADPH-dependent activity. Growth of the library and controls was monitored throughout the selection. As expected, the negative controls (APEQS transformed with pBbA5c_MEV_da or pBbA5c_empty) showed no detectable growth over this time course, whereas the positive control (APEQS transformed with pBbA5c_MEV_sa) grew robustly to a high final OD (Fig. 3b). The library exhibited a longer lag phase than the positive control but ultimately grew robustly over the 72-hour time course (Fig. 3b). This pattern is consistent with rare NADPH-specific, high-activity variants initially present at low frequency becoming enriched over time and eventually dominating the culture. To isolate the fitness effect of activity-enhancing plasmid-borne mutations from potential genomic mutations circumventing selective pressure, the plasmids from each replicate evolution were isolated from the cultures after outgrowth. The isolated plasmids were then used to transform fresh APEQS cells and individual 84 clones were isolated on non-selective media and then inoculated into the selective condition to determine growth rate relative to APEQS_MEV_da and APEQS_MEV_sa. The growth rate of 84 clones was quantified (Fig. 3c).

### 2.4. Evaluation of HMGR variants with altered cofactor specificity

The 20 fastest growing clones of APEQS cells transformed with evolved library DNA were then sequenced (Fig. 4a). To clearly visualize patterns in the enrichment of specific amino acids at the three positions the sequences were used to generate a sequence logo (Fig. 4b). This logo revealed high conservation of either serine or asparagine at position 148 and arginine at position 152, which are also found in HMGR_sa (Fig. 4a). Structural analysis indicated that the long, polar Arg152 side chain stabilizes the 2′-phosphate via hydrogen bonding, and that introducing a polar residue at position 148 (e.g., Asn148) provides additional polar contacts that further stabilize the 2′-phosphate (Supplementary Fig. 2b). There was limited consensus of amino acid identity of position 146. However, none of the sequenced clones contained a negatively charged amino acid at 146. Although many residues are tolerated at position 146, it appears the wild-type negative charge must be altered to prevent electrostatic repulsion with NADPH’s 2′-phosphate, which lies close to residue 156 (Supplementary Fig. 2a,b)

**Figure 4.**
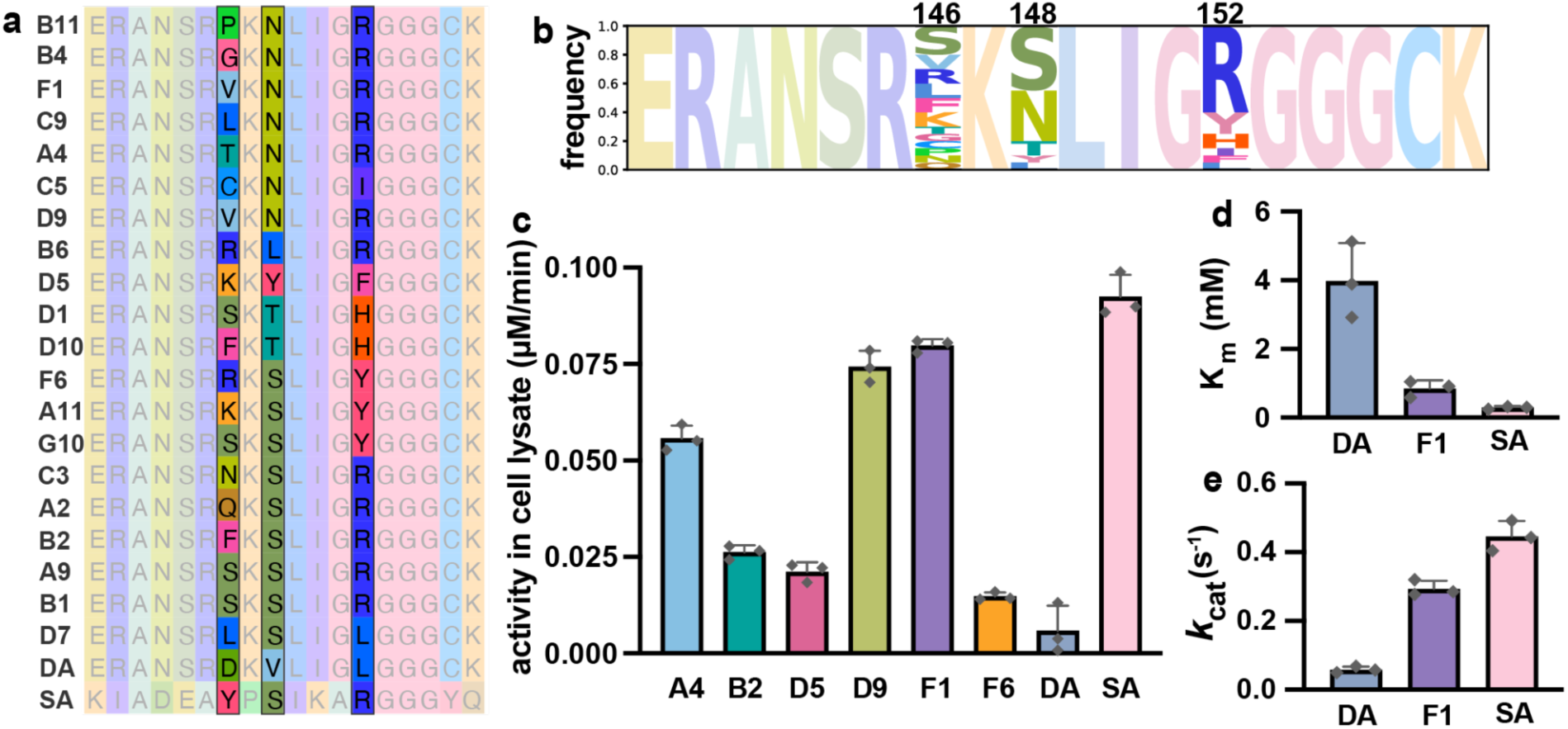
Analysis of HMGR_da variants a, Sequence alignment of mutant HMGR_da variants as well as WT HMGR_da and HMGR_sa. **b,** Logo motif of mutant HMGR_da sequences. **c,** HMGR activity in cell lysates containing the variants of interest was measured in the reverse direction by monitoring NADPH generation by absorbance at 340 nm in a reaction mixture containing 2 mM free CoA, 2 mM mevalonate and 5 mM NADP+. **d,** K_m_ of HMGR_da (4.0 ± 1.1 mM), HMGR_sa (0.29 ± 0.04 mM) and HMGR_da_F1 (0.85 ± 0.23 mM) for NADP^+^. **e,** *k*_cat_ of HMGR_da (0.058 ± 0.009 s^-1^), HMGR_sa (0.45 ± 0.04 s^-1^) and HMGR_da_F1 (0.29 ± 0.02 s^-1^) in the HMG-CoA forming direction.

To further characterize the fastest-growing mutants, we assayed the NADP^+^-reducing activity of the HMGR variants in cell lysates (Fig. 4c). We found that the F1 variant exhibited the highest NADP^+^-reducing activity in a reaction mixture also containing mevalonate and free-CoA. To quantify and compare the kinetic parameters of the F1 variant, we purified HMGR_da, HMGR_sa and HMGR_da_F1 and assayed the *K*_m_ and *k_cat_* of each enzyme. We were unable to saturate the enzyme in the forward, NADPH-oxidizing direction and therefore assayed the enzymes in the reverse direction to determine the *k_cat_* and the *K*_m_ for NADP^+^.

HMGR_da_F1 exhibited a 4.7-fold decrease in apparent *K*_m_ for NADP^+^ and 5.0-fold increase in *k_cat_*, corresponding to a 23.4-fold increase in NADP^+^-dependent catalytic efficiency (*k_cat_/K_m_*) relative to HMGR_da. These results indicate that the described selection scheme, supported by an acetate co-feeding strategy, provides sufficient evolutionary pressure to enable directed evolution. Despite not supplying the direct substrate of HMGR, we were able to resolve stoichiometric constraints precluding effective growth coupling by feeding acetate in a manner that could be widely generalized to many partially-reduced acetyl-CoA-derived bioproducts.

### 2.5. Growth coupling of alternative acyl-CoA substrates

To demonstrate that this growth coupling scheme involving the co-feeding of a secondary, non-utilizable carbon source is generalizable beyond acetyl-CoA, we chose to growth-couple the reduction of propionate in a similar manner to acetate. We generated a strain that can activate exogenously supplied propionate to propionyl-CoA without the ability to catabolize propionyl-CoA (Fig. 5a). We accomplished this by an in-frame deletion of the gene *prpC,* which encodes 2-methylcitrate synthase, a key enzyme in the methylcitrate cycle used to catabolize propionate (Fig. 5a).^50^ Our in-frame deletion preserved the open reading frames of downstream genes in the *prpBCDE* operon and crucially the native PrpE enzyme responsible for activation of propionate to propionyl-CoA.^26,51^ This MX203 strain with *prpC* deletion was named PPEQS and used for subsequent growth coupling of propionyl-CoA reduction. To partially reduce propionyl-CoA in a NADPH-dependent manner, we again used the enzyme *Rp*PduP-NP.^43^

**Figure 5.**
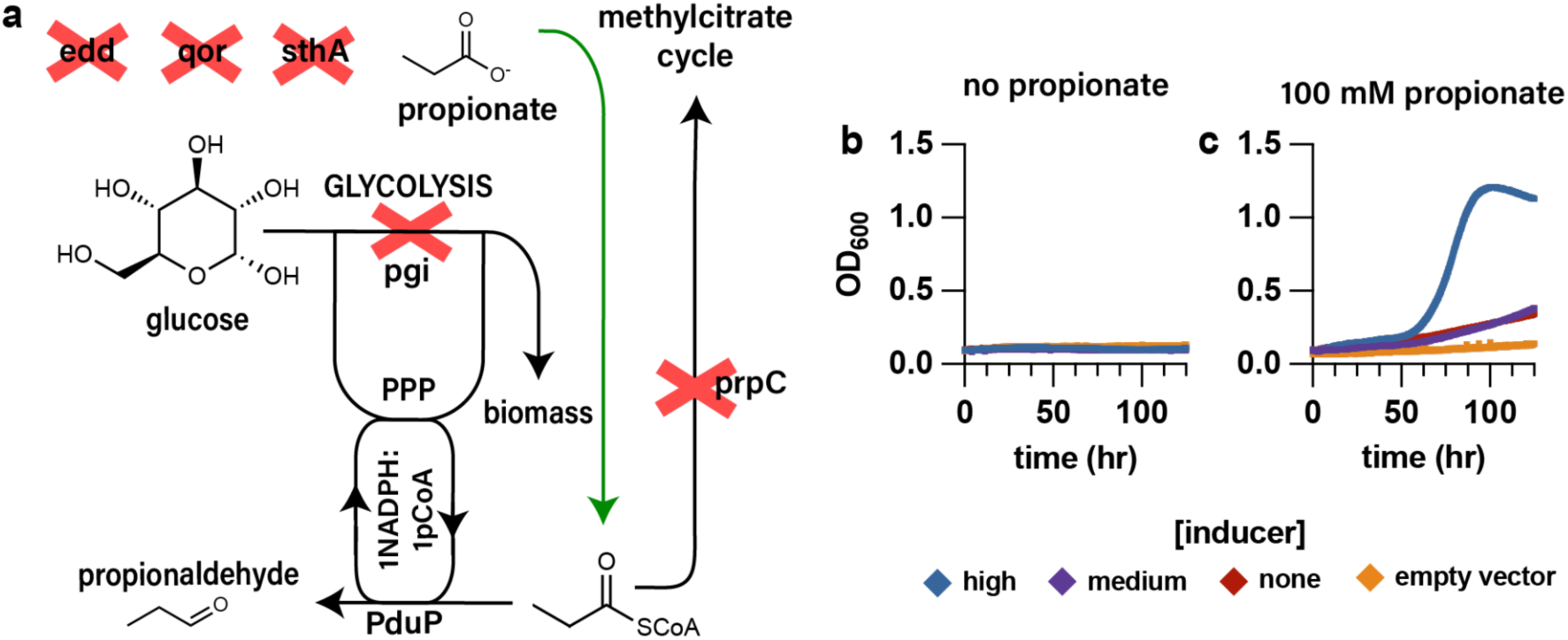
Growth coupling of the partial reduction of propionyl-CoA by propionate co-feeding a, Schematic diagram showing knockouts and carbon fluxes in PPEQS. Propionate is supplied as a secondary carbon source to generate propionyl-CoA which is partially reduced by *Rp*PduP-NP. Deletion of *prpC* prevents use of propionate as a sole carbon source. **b,** Expression of *Rp*PduP-NP in strain PPEQS_PduP is not sufficient to rescue growth without supplementation of propionate. **c,** Expression of *Rp*PduP-NP in strain PPEQS_PduP successfully rescues growth when 100 mM propionate is supplemented to the medium.

Plasmid pQE_PduP containing *Rp*PduP-NP^43^ was used to transform PPEQS generating strain PPEQS_PduP. When no propionate was added as a secondary carbon source, PPEQS_PduP did not grow over the assayed time-course (Fig. 5b). Using 100 mM propionate as a secondary carbon source, we were able to rescue the growth of PPEQS_PduP when the expression of *Rp*PduP-NP was induced with IPTG (Fig. 5c). Induction-dependent propanol production was verified to ensure that propionaldehyde was being produced and detoxified to propanol as expected (Supplementary Fig. 3).

These results indicate that a propionate co-feeding strategy, like the acetate co-feeding strategy described above, satisfied the stoichiometric constraints that otherwise preclude the rescue of PPEQS_PduP by *Rp*PduP-NP. These results bear special importance for the growth coupling of partially reduced polyketides. Many polyketide synthases use methylmalonyl-CoA, derived from the carboxylation of propionyl-CoA, as extender units that are subsequently reduced by downstream reductase domains.^26,52^ The growth coupling of the partial reduction of propionyl-CoA sets a conceptual framework for the growth coupling of reducing polyketide synthases by the described redox-balance principles.

## 3. Discussion and Conclusion

Maintenance of redox balance is a fundamental principle in biology, and recent work has leveraged this principle to build growth-coupled selections for enzyme evolution. Here, we define stoichiometric constraints that have limited the scope of directed evolution through redox-balance growth coupling. In prior implementations, these constraints were typically addressed by feeding a direct precursor, restricting growth-coupled approaches to reactions whose immediate substrates can be supplied exogenously. In contrast, we expand redox-based growth coupling to multistep pathways using a versatile co-feeding strategy. A key innovation is the use of acetate as a redox-neutral carbon source that supports bioproduction while decoupling carbon availability from NADPH generation.

Using this framework, we show how selective pressure can be applied to pathways involving iterative acyl-chain elongation for the purpose of directed evolution. This class of chemistry underlies many pathways of industrial interest including but not limited to fatty acid and polyketide biosynthesis.^8^ However, we also acknowledge limitations of our described selection scheme to help clarify appropriate use cases. To this end we describe a dynamic range of selective pressure applied through the growth-coupled scheme, highlighting an often-overlooked limitation of growth-coupled selective systems: they require a baseline flux through the engineered pathway to produce a measurable fitness advantage.

To demonstrate a suitable directed-evolution application, we altered the cofactor specificity of an NADH-dependent HMGR to accept NADPH by applying selection to a saturation-mutagenesis library expressed in APEQS. Alteration of enzyme cofactor specificity is a key tool for modulating pathway kinetics and thermodynamics^53^ and has been used in metabolic engineering efforts to increase product titer^54,55^ or accomplish other engineering goals^25^. Broadly, the platform should be applicable to other enzyme-engineering objectives (e.g., substrate specificity or catalytic efficiency), provided improved variants increase redox flux sufficiently to yield a growth advantage within the selection’s dynamic range.

Finally, we explored substrates other than acetyl-CoA to test whether this selective pressure generalizes to other acyl-CoAs. We showed that NADPH-dependent propionaldehyde production can be growth-coupled in the same manner as acetaldehyde production, supporting the broader applicability of the scheme. Notably, requiring the pathway to utilize a secondary carbon source creates a route to selection for substrate specificity: promiscuous enzymes can be pressured to accept the supplemented substrate when it is necessary to satisfy the stoichiometric constraints of growth coupling. Designing systems in which the target enzyme must process the supplemented carbon source may therefore provide a general strategy for evolving substrate specificity in future work.

This work sets a conceptual framework for the growth coupling of the diverse pathways involving acyl-CoA extension and reduction and seeks to support future work enabling the directed evolution of metabolic pathway enzymes for efficient chemical production.

## 4. Methods

### 4.1. Cell Culture

For routine culturing, bacterial strains were grown in 2xYT medium (16 g/L tryptone, 10 g/L yeast extract, and 5 g/L sodium chloride), supplemented with the appropriate antibiotic based on the plasmid(s) present in the strain: carbenicillin (100 µg/mL), kanamycin (50 µg/mL), or chloramphenicol (25 µg/mL). For growth coupling experiments and strain evolution, bacterial strains were grown in a minimal MOPS medium (Teknova M2106) containing 2% glucose (w/v) supplemented with the appropriate antibiotic based on the plasmid(s) present in the strain.

### 4.2. DNA Cloning and Assembly

For the construction of pQE_PduP the codon-optimized DNA sequence for *Rp*PduP was synthesized by Integrated DNA Technologies and inserted into the pQE backbone using Gibson assembly. Sequence-verified plasmid containing the WT *Rp*PduP was then used as a template to introduce mutations P164G, I199R, N203L, and I222L by amplification with mutagenic primers and reconstitution of the final plasmid using Gibson assembly. The ADH1 gene from *Saccharomyces cerevisiae* was then amplified from genomic DNA and inserted into the plasmid containing the mutant *Rp*PduP using NEB HiFi assembly to generate the final pQE_PduP plasmid.

Plasmid pBbA5c_3-HB was constructed by amplification of plasmid pBbA4c_HMGR_sa (registry # JBEI_17081) ^48^ to retain only the backbone and *atoB* gene, the *phaB* gene from *Cupriavidus necator* was amplified from genomic DNA and these fragments were subsequently assembled using NEB HiFi Master Mix.

Constructs for protein purification were generated by insertion of *hmgr* genes into the pET28a backbone for T7-based expression. *hmgr* genes and pET28a backbone were amplified via PCR and then assembled via NEB HiFi Assembly.

All DNA constructs were verified via whole plasmid sequencing using Oxford Nanopore technology (Plasmidsaurus).

### 4.3. Transformation

Plasmid DNA was introduced into cells via electroporation. Before transformation, log-phase *E. coli* cultures were pelleted at 4000 × g, the supernatant was discarded, and the cells were washed three times with 10% glycerol. For each transformation, 1 µL of plasmid DNA (100 ng/µL) was mixed with 25 µL of washed cells and transferred to a 0.1 cm electroporation cuvette. Electroporation was carried out at 1.55 kV. Immediately following electroporation, 1 mL of 2xYT medium was added to the cuvette contents and transferred to a 1.75 mL Eppendorf tube. Cells were recovered at 37 °C for 1 hour, then plated onto LB agar plates containing the appropriate antibiotic for selection.

### 4.4. Genome Engineering

Gene knockouts were performed by lambda red recombineering. Plasmid pKD4 containing a kanamycin resistance cassette was amplified by PCR using primers containing overlaps homologous to the chromosomal target. Allelic exchange was facilitated by the expression of the Red Disruption System from plasmid pKD46.^56^ To remove the kanamycin cassette from MX203 *prpC::kan*, plasmid pCP20 was used.

Removal of the kanamycin cassette was important in this case to maintain the expression of downstream CoA-ligase *prpE*.^56^ The kanamycin cassette was retained in strain APEQS due to concerns about crossover between the FRT (Flp recombination target) scar at the Δ*pgi* site and FRT sites introduced at the neighboring *aceA* gene locus, which could cause a loop-out event, removing the intervening genes.

### 4.5. Bacterial Growth Measurements

Strains were cultured in 2xYT medium overnight at 30 °C. The next day, dense overnight cultures were diluted into fresh 2xYT at a ratio of 1:50 and grown for approximately 2 hours at 37 °C until the optical density of the cultures (OD600) had reached approximately 0.4, at which point 0.5 mM IPTG was added and the temperature was reduced to 30 °C. Cultures were incubated for another 2 hours to enable expression of pathway genes. After 2 hours, the cells were washed three times in

MOPS medium containing 2% glucose and then diluted to an OD of 0.1 in a selective medium containing the indicated supplementary carbon source and inducer. For genes expressed from the medium-copy pBbA5c backbone (mevalonate and 3-HB production pathways), concentrations of IPTG used were 0.5 and 0.05 mM for high and medium expression respectively. For genes expressed on the high-copy pQE backbone 0.05 and 0.005 mM IPTG were used for high and medium expression respectively. 0.5 mL of the diluted culture in a minimal medium containing the appropriate antibiotics and inducer was transferred to a clear-bottom 48-well plate (Falcon) and incubated in a Molecular Devices M2 plate reader with shaking at 30 °C for the indicated amount of time. Optical density at 600 nm was measured every 10 minutes over the time course. Raw optical density measurements were adjusted to reflect a standard 1 cm path length. Doubling times were calculated according to the method described in He et al^57^ and were converted to growth rate using the function µ=ln(2)/T_d_.

### 4.6. Quantification of bioproduct formation

Quantification of 3-HB and propanol was performed using high-performance liquid chromatography with refractive index detection (HPLC-RID). An Agilent Series 1200 was used to deliver a 50 µL injection volume of clarified culture supernatant, arising from a growth experiment, to an Aminex HPX-87H column (Bio-Rad) at 65 °C with subsequent detection by RID maintained at 50 °C. Separation and quantification were performed in a 5 mM H_2_SO_4_ solution flowing at a rate of 0.600 mL/min.

Quantification of mevalonate was performed by liquid chromatography–mass spectrometry (LC–MS) due to overlapping peaks arising from the MOPS culture medium at the retention time of mevalonate. An Agilent 1260 Infinity II LC system equipped with a MSD/iQ mass spectrometer was used to deliver a 10 µL injection volume of clarified culture supernatant to an Astec^®^ C18 HPLC Column (5 µm particle size, L × I.D. 15 cm × 4.6 mm) (Supelco) for separation at room temperature with subsequent detection by MSD/iQ mass spectrometer. Ultrapure water and methanol (MeOH) containing 0.1% formic acid were used as the mobile phase in the following gradient: 5% MeOH to 85% MeOH over 4 min, 85% MeOH to 98% MeOH over 3.8 min, both at a flow rate of 0.420 mL/min, then 98% MeOH to 5% MeOH over 0.4 min and hold at 5% MeOH for 4.8 min at a flow rate of 0.650 mL/min. Mevalonate was monitored by a negative mode single-ion monitoring (SIM) at m/z = −147.

### 4.7. Library Construction and Growth-Coupled Evolution

Plasmid pBbA5c_MEV_da was used as the template for library generation.

Primers were designed containing NNK codons at the sites complimentary to codons 146, 148, and 152 as well as BsaI restriction sites (forward primer: 5’-cacaccaGGTCTCagccgcNNKaaaNNKctgatcggtNNKggcggtggctgtaaagatattga-3’; reverse primer 5’-cacaccaGGTCTCgcggctatttgcgcgttcaatgat-3’). BsaI restriction sites were used for the scarless recircularization of the plasmid pBbA5c_MEV_da now containing the NNK codons, generating DNA library pBbA5c_MEV_da_lib. This library was transformed into XL1-Blue and the number of colony-forming units (CFU) arising from transformation was determined by plating serial dilutions on selective agar medium. We obtained 9.2×10^5^ transformants (115-fold oversampling of the 8.0×10^3^ theoretical variants), corresponding to an expected 99.999% variant representation under a uniform distribution assumption. The XL1-Blue transformants were scraped from their selective medium and pooled. The library was then extracted from the pooled transformants by miniprep (QIAGEN) to generate purified, pooled library DNA.

The purified library in triplicate, as well as control DNA, was then transformed into APEQS following the method described above. After recovery, 10 µL of each sample was serially diluted to estimate the CFU resulting from transformation. Each transformation resulted in over 10^6^ transformants, ensuring library coverage was maintained. The remaining cells in the recovery medium were then diluted onto 5 mL of 2xYT containing chloramphenicol and kanamycin and incubated at 37 °C until the OD600 reached 0.6, at which point 0.5 mM IPTG was added to induce gene expression. The cultures were grown for another 4 hours until OD600 was approximately 1.5. Once the cultures were sufficiently grown they were harvested and washed three times with minimal MOPS medium. The washed cells were then inoculated into a selective medium (MOPS medium containing chloramphenicol and kanamycin, 2% glucose, 0.5 mM IPTG and 100 mM acetate) at an OD of 0.175. The selection was conducted for 72 hours, at which point individual clones were isolated from the evolved library for further analysis.

### 4.8. *In vitro* biochemical analysis using cell lysates

Overnight cultures of APEQS_MEV_sa, APEQS_MEV_da, and APEQS transformed with library variants of interest were inoculated at 1:20 dilution into 2xYT and grown at 37 °C. Once cultures had reached an OD_600_ of ∼0.4, they were induced with 0.5 mM IPTG and grown overnight at 30 °C to express HMGR proteins. 5 mL of cell culture was harvested by centrifugation and resuspended in 500 µL of lysis buffer (50 mM Tris-HCl pH 7.4, 300 mM NaCl, 0.3 mg/mL lysozyme, 1 mM DTT). Upon suspension, 500 µL of soda-lime glass microspheres were added as a mechanical lysing agent. These mixtures were transferred to a Retsch MM400 mixer mill and milled at 30 Hz for 10 minutes. The mixture was then clarified by centrifugation at 10,000 × g for 10 minutes at 4 °C. The cell lysate was isolated and stored at −80 °C for use in kinetic assays.

To compare the activity of generated HMGR mutants to that of HMGR_da and HMGR_sa, an *in vitro* assay was conducted using the clarified cell lysate. Standard reaction conditions were 50 mM Tris-HCl at pH 8.5, 2 mM mevalonate, 2 mM free CoA, and 5 mM NADP+ at 30 °C. Reactions were conducted in 100 µL volumes and initiated by the addition of 5 µL of cell lysate. Specific activity was determined by measuring Abs_340_ to monitor NADPH generation in a Nunc 96-well optical-bottom microplate (Thermo Fisher Scientific) using a Molecular Devices M2 Spectrophotometer.

### 4.9. Protein purification

Cultures of BL21 (DE3) *E. coli* strains harboring pET28a_HMGR_da, pET28a_HMGR_da_F1 and pET28a_HMGR_sa were inoculated into 500 mL of TB with kanamycin at an OD600 of 0.1 and grown at 37 °C with 200 rpm shaking. Upon reaching OD_600_ ∼0.4, cultures were placed on ice for 10 minutes prior to addition of IPTG to a final concentration of 0.1 mM. Cultures were then grown overnight at 18 °C for overexpression of HMGR proteins.

After 18 hours of overexpression, the cells were harvested and lysed by sonication in 30 mL of lysis buffer (50 mM Tris-HCl pH 7.4, 300 mM NaCl, 10% (v/v) glycerol, 5 mM imidazole). Post-lysis, the cells were centrifuged at 5,000 × g for 15 minutes at 4 °C. Clarified lysates were isolated for purification. 6 mL of Ni-NTA resin beads (Thermo Fisher Scientific) suspended in lysis buffer were added to the resulting lysates and equilibrated by rotation at 4 °C for 1 hr. The equilibrated lysate-resin mixtures were loaded onto gravity flow columns to remove lysis buffer from the resin beads. Non-His-tagged proteins were removed from the resin bed with 6 mL of wash buffer (50 mM

Tris-HCl pH 7.4, 300 mM NaCl, 40 mM imidazole) by washing 4 times. His-tagged proteins were removed from Ni-NTA beads with elution buffer (50 mM Tris-HCl pH 7.4, 300 mM NaCl, 10% (v/v) glycerol, 200 mM imidazole) in 2 mL fractions. Purified fractions were dialyzed to remove imidazole and flash frozen in protein storage buffer (50 mM Tris-HCl pH 7.4, 300 mM NaCl, 10% (v/v) glycerol).

### 4.10. *In vitro* biochemical analysis using purified protein

HMGR enzymatic activity was determined in the mevalonate-oxidizing direction due to our inability to saturate the enzyme in the forward direction. An enzyme concentration of 20 ng/µL was used. Standard reaction conditions were 100 mM Tris at pH 8.5, 50 mM NaCl, 1.5 mM mevalonate, 1.5 mM CoA, and various concentrations of NADP^+^ (HMGR_sa: 0.1 mM, 0.25 mM, 0.5 mM, 2 mM and 8 mM; HMGR_da and _F1: 0.25 mM, 0.5 mM, 2 mM, 5 mM, 8 mM and 20 mM) in 100 µL reactions. Kinetic parameters were determined by measuring A340 to monitor NADPH generation in a Nunc 96-well optical-bottom microplate (Thermo Fisher Scientific) using a Molecular Devices M2 Spectrophotometer held at 30 °C.

## Acknowledgments

This work was supported by the US Department of Energy (DOE) Joint BioEnergy Institute (https://www.jbei.org), supported by DOE, Office of Science, Biological and Environmental Research Program, under contract DE-AC02-05CH11231 between DOE and Lawrence Berkeley National Laboratory (J.D.K.), the DOE Distinguished Scientist Fellow Program (J.D.K.), the Philomathia Foundation (J.D.K.), National Institutes of Health grant R01 AT010593-02 (J.D.K.), and National Science Foundation grant 2036849 (J.D.K.).

## Author statement

Conceptualization: A.E.C.; Data Curation: A.E.C.; Formal Analysis: A.E.C.; Methodology: A.E.C.; Investigation: A.E.C., B.C., M.H., S.P., M.G., A.S., S.S., J.G., Y.C.; Software: A.E.C. and Y.C.; Visualization: A.E.C.; Validation: A.E.C.; Writing - original draft: A.E.C.; Funding Acquisition, Resources, and Supervision: C.J.P., and J.D.K.; Writing - review and editing: All authors.

## Data Availability

Raw proteomic data have been deposited to the PRIDE database with accession code PXD075294 (reviewer token P6iPkUfJrsRI).

