## Supplementary Materials for "Expanding the scope of redox-balance growth coupling techniques with a carbon cofeeding strategy"

### Supplementary Table 1: Strains used in this study

| Strain name | Genotype | Description | Registry # * | Reference |
|---|---|---|---|---|
| Mx203 | *E. coli* BW25113 Δ*pgi* Δ*edd* Δ*qor* Δ*sthA* | *E. coli* strain containing knockouts of genes to arrest growth due to NADPH redox imbalance when grown with glucose as the sole carbon source | JPUB_xx | [1] |
| APEQS | Mx203 *aceA::kan* | Mx203 containing an additional deletion in *aceA* to prevent growth on acetate as a sole carbon source. APEQS acronym denotes the first letter of each of the five gene deletions relative to WT *E. coli* | JPUB_xx | This study |
| APEQS_PduP | APEQS with plasmid **pQE_PduP** | APEQS transformed with plasmid **pQE_PduP** for the NADPH-dependent production of acetaldehyde and NADH-dependent detoxification to ethanol | JPUB_xx | This study |
| APEQS_3-HB | APEQS with plasmid **pBbA5c_3-HB** | APEQS transformed with plasmid **pBbA5c_3-HB** for the NADPH-dependent production of (R)-3-hydroxybutyrate | JPUB_xx | This study |
| APEQS_MEV_sa | APEQS with plasmid **pBbA5c_MEV_sa** | APEQS transformed with plasmid **pBbA5c_MEV_sa** for the NADPH-dependent production of mevalonate | JPUB_xx | This study |
| APEQS_MEV_da | APEQS with plasmid **pBbA5c_MEV_da** | APEQS transformed with plasmid **pBbA5c_MEV_da** for the NADH-dependent production of mevalonate | JPUB_xx | This study |
| PPEQS | Mx203 Δ*prpC* | Mx203 containing an additional deletion in *prpC* to prevent propionate catabolism. PPEQS acronym denotes the first letter of each of the five gene deletions relative to WT *E. coli* | JPUB_xx | This study |
| PPEQS_PduP | PPEQS with plasmid **pQE_PduP** | PPEQS transformed with plasmid **pQE_PduP** for the NADPH-dependent production of propionaldehyde and subsequent detoxification to propanol | JPUB_xx | This study |
| BL21(de3)_HMGR_sa | BL21(de3) with plasmid **pET28a_HMGR_sa** | BL21(de3) transformed with plasmid **pET28a_HMGR_sa** for the purposes of protein purification | JPUB_xx | This study |
| BL21(de3)_HMGR_da | BL21(de3) with plasmid **pET28a_HMGR_da** | BL21(de3) transformed with plasmid **pET28a_HMGR_da** for the purposes of protein purification | JPUB_xx | This study |
| BL21(de3)_HMGR_F1 | BL21(de3) with plasmid **pET28a_HMGR_da_F1** | BL21(de3) transformed with plasmid **pET28a_HMGR_da_F1** for the purposes of protein purification | JPUB_xx | This study |
| XL1-Blue | *E. coli* XL1-Blue (*rec*A1 *end*A1 *gyr*A96 *thi*-1 *hsd*R17 *sup*E44 *rel*A1 lac [F´ *proAB lacI*$^q$Z*ΔM15* Tn*10* (Tet r )]) | Used for DNA storage and preparation | JPUB_xx | Commercially available (Agilent) |
| BL21(de3) | *E. coli* BL21(de3) (*fhuA2 [lon] ompT gal (λ DE3) [dcm]* Δ*hsdS λ DE3 = λ sBamHIo* Δ*EcoRI-B int::(lacI::PlacUV5))::T7 gene1) i21* Δ*nin5* | Used for T7-based protein expression and purification | JPUB_xx | Commercially available (Agilent) |

Plasmid names are bolded  * registry information will be added prior to final submission to ensure consecutive numbering

### Supplementary Table 2: Plasmids used in this study

| Plasmid name | Description | Registry # * | Reference |
|---|---|---|---|
| **pQE_PduP** | Expression construct for *Rhodopseudomonas palustris* propanediol utilization protein (*Rp*PduP) with mutations conferring NADPH specificity (P164G, I199R, N203L, and I222L).[2] Additionally, this construct contains NADH-dependent ADH1[3] from *Saccharomyces cerevisiae* to supplement native aldehyde dehydrogenase and ensure that accumulation of aldehydes does not inhibit growth during growth coupling. | JPUB_xx | This study |
| **pBbA5c_3-HB** | Expression construct for *E. coli* thiolase AtoB and *Cupriavidus necator* PhaB for the production of 3-HB. | JPUB_xx | This study |
| **pBbA5c_MEV_sa** | Expression construct for *E. coli* thiolase AtoB, *Cupriavidus necator* PhaB and HMGR from *Staphylococcus aureus* for the production of mevalonate. | JPUB_xx | [4] |
| **pBbA5c_MEV_da** | Expression construct for *E. coli* thiolase AtoB, *Cupriavidus necator* PhaB and HMGR from *Delftia acidovorans* for the production of mevalonate. | JPUB_xx | [4] |
| **pET28a_HMGR_sa** | Protein overexpression construct for HMGR from *Staphylococcus aureus* for the purpose of protein purification. | JPUB_xx | This study |
| **pET28a_HMGR_da** | Protein overexpression construct for HMGR from *Delftia acidovorans* for the purpose of protein purification. | JPUB_xx | This study |
| **pET28a_HMGR_da_F1** | Protein overexpression construct for HMGR from *Delftia acidovorans* with mutations D146V, V148N and L152R for protein purification. | JPUB_xx | This study |

* registry information will be added prior to final submission to ensure consecutive numbering

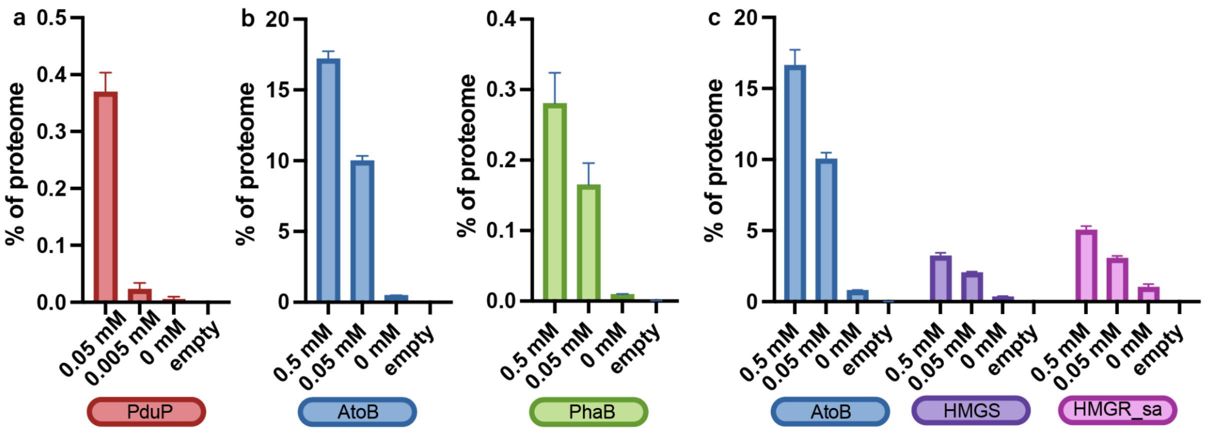


**Supplementary Figure 1 Proteome allocation to pathway genes**

**a,** The percentage of the proteome allocated to *Rp*PduP-NP under growth coupled conditions. **b,** The percentage of the proteome allocated to enzymes supporting 3-HB production from acetyl-CoA, namely AtoB (blue) and PhaB (green) expressed from plasmid pBbA5c_3-HB under growth-coupled conditions in APEQS. **c,** Percentage of the proteome allocated to enzymes supporting the production of mevalonate from acetyl-CoA, namely AtoB (blue), HMGS (purple) and HMGR (pink) from plasmid pBbA5c_MEV_sa under growth coupled conditions.

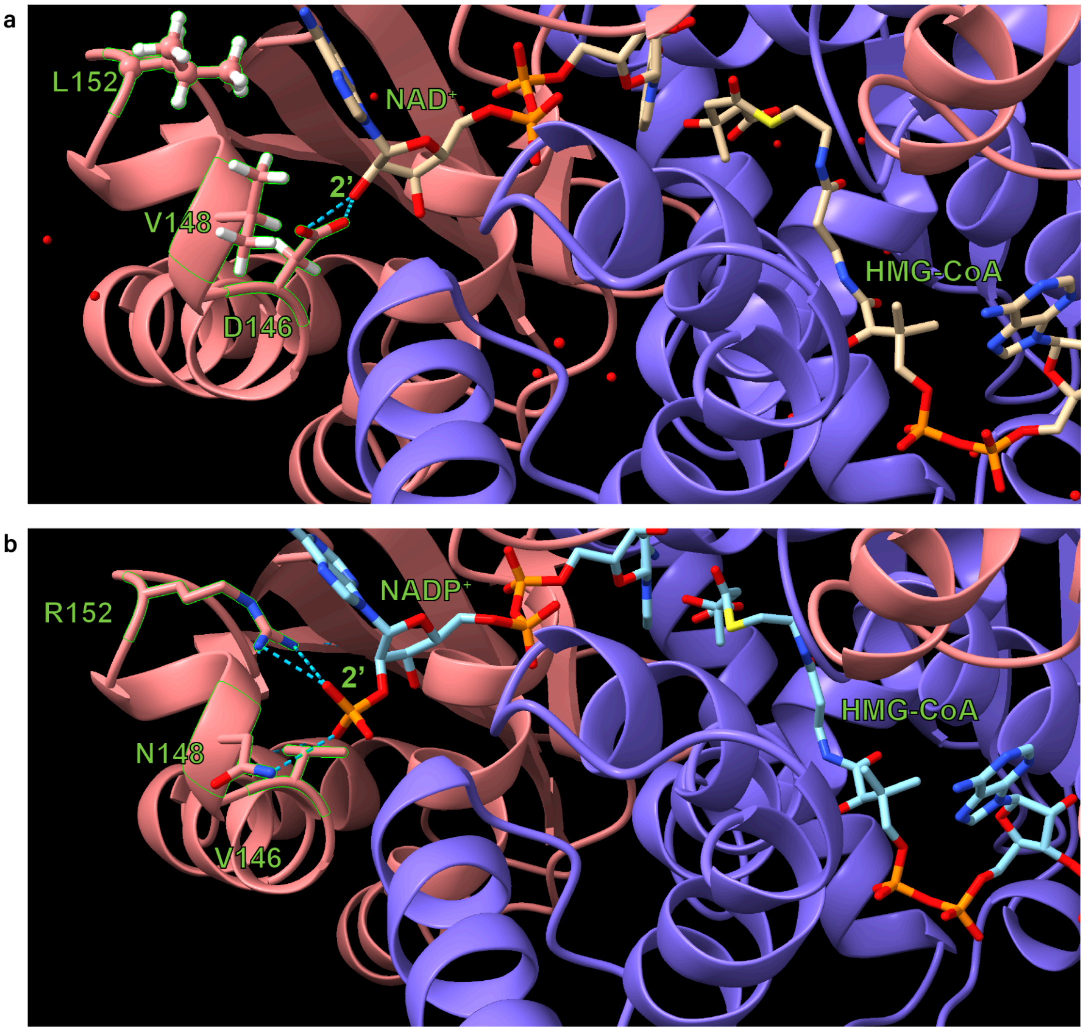


**Supplementary Figure 2 Structural analysis of HMGR substrate specificity**
**a,** The predicted structure of the HMGR_da dimer was generated using Boltz-2x and is shown (chain 1: pink, chain 2: purple) with substrate HMG-CoA and the native cofactor $NAD^+$. Substrate and cofactors are oriented within the predicted structure of HMGR_da based on alignment with PDB: 1QAX (Pseudomonas mevalonii HMGR with NAD+ and HMG-CoA). Key residues targeted for saturation mutagenesis (D146, V148, and L152) are labeled, showing proximity to the 2' hydroxyl group that is phosphorylated in NADPH. Hydrogen bonds formed between the residues of interest and the 2' position of the nicotinamide cofactor are denoted with blue dashed lines. **b,** The predicted structure of HMGR_da_F1 (HMGR_da D146V, V148N and L152R) with substrate HMG-CoA and cofactor $NADP^+$ was generated using Boltz-2x and is shown in its dimer form (chain 1: pink, chain 2: purple). Key residues in HMGR_da_F1 mutated relative to HMGR_da are labeled. Hydrogen bonds formed between mutant residues and the 2' phosphate of $NADP^+$ are indicated by dashed blue lines.

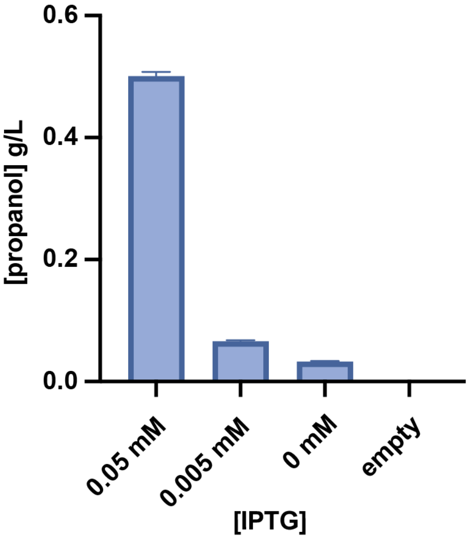


**Supplementary Figure 3 Propanol production in PPEQS_PduP strain**
Propanol production was assayed from strain PPEQS_PduP after growth in minimal medium containing 2% glucose and 100 mM propionate with the specified concentrations of IPTG. Results indicate that plasmid pQE_PduP enabled the production of propanol in PPEQS_PduP in a minimal medium containing 100 mM propionate.
